# Incorporating polygenic scores in the twin model to estimate genotype-environment covariance: exploration of statistical power

**DOI:** 10.1101/702738

**Authors:** Conor V. Dolan, Roel C. A. Huijskens, Camelia C. Minică, Michael C. Neale, Dorret I. Boomsma

## Abstract

The assumption in the twin model that genotypic and environmental variables are uncorrelated is primarily made to ensure parameter identification, not because researchers necessarily think that these variables are uncorrelated. Although the biasing effects of such correlations are well understood, it would be useful to be able to estimate these parameters in the twin model. Here we consider the possibility of relaxing this assumption by adding polygenic score to the (univariate) twin model. We demonstrated numerically and analytically this extension renders the additive genetic (A) – unshared environmental correlation (E) and the additive genetic (A) - shared environmental (C) correlations simultaneously identified. We studied the statistical power to detect A-C and A-E correlations in the ACE model, and to detect A-E correlation in the AE model. The results showed that the power to detect these covariance terms, given 1000 MZ and 1000 DZ twin pairs (α=0.05), depends greatly on the parameter settings of the model. We show fixing the estimated percentage of variance in the outcome trait that is due to the polygenic scores greatly increases statistical power.

---

The classical twin design (Eaves, et al. 1978; Martin et al. 1997) arguably has been one of the most productive genetically informative designs in the study of human traits (Polderman et al. 2015). Twin studies have contributed greatly to our present knowledge concerning genetic and environmental contributions to individual differences in psychological traits (e.g., Plomin, Defries, Knopik, and Neiderhise, 2016; Turkheimer, 2002; Plomin and Deary, 2016) and multivariate and longitudinal extensions of the classical twin design (Martin and Eaves, 1977) have provided insights into the etiology of comorbidity and stability of traits and disorders. However, the interpretation of results from the uni- or multivariate model depend on its model assumptions. The main assumptions of the CTD that are often evoked concern genotype-environment covariance (assumed to be absent), genotype-environment interaction (assumed to be absent), the equal environment assumption (the influence of shared environment assumed to be equal in MZ and DZ twins), and parental mating (assumed to be random for the trait that is analyzed). Given these assumptions, the results from the CTD can provide unbiased estimates of additive genetic (A), unshared environmental (E), and shared environmental (C) <u>or</u> dominance (D) variance components. The effect of violations of these are well understood (Verhulst & Hatemi, 2013; Purcell, 2002; Keller et al. 2009), so that estimates of variance components obtained in the twin model may be interpreted in the light of possible model violations. For instance, a significant estimate for C in the CTD may the results of assortative mating, or of positive A-C correlation. A correlation between A and E contributes to the A variance component.

Many papers have been devoted to the detection and accommodation of model violations, either within the CTD (e.g., Purcell, 2002, Molenaar, et al. 2012, Eaves & Erkanli, 2003, Carey, 1986; Dolan, et al. 2014; Beam & Turkheimer, 2013), or in extended designs (e.g., Plomin, Loehlin, and DeFries, 1985; Narusyte, et al, 2008; Fulker, 1988; D’Onofrio, et al, 2003; Keller et al, 2009; Heath et al., 1985, Maes, et al, 2006). The aim of the present paper is to demonstrate that incorporating polygenic scores in the classical twin design in principle allows us to estimate the covariance between A and C and between A and E. In developmental psychology, such covariance terms are viewed as plausible, stemming from processes giving rise to passive, active or evocative genotype-environment covariance (Plomin, DeFries, and McClearn, 1977; Scarr and McCartney, 1983). Knafo & Jaffee, 2013; Kendler, 2012; Rutter and Silberg, 2002). Beyond demonstrating the principle, we explore the statistical power to estimate rGE under alternative models.

The idea of incorporating measured genetic information, either genetic variants or polygenic scores, in the twin design is not new. Previous approaches concerned measured genetic markers in non-parametric linkage analysis (e.g., Haseman and Elston, 1977; Nance and Neale, 1989) or combined linkage-association analysis (e.g., Fulker, Cherny, Sham, Hewitt, 1999). Van den Oord and Snieder (2002) presented an extended twin model with measured genetic variable to test association in the presence of stratification and to test causal relationships. In a longitudinal study of attention problems, van Beijsterveldt et al. (2012) incorporated measured candidate gene information in a common factor twin model to test the association with the latent phenotype attention problems. Minică et al. (2018) presented an integration of the CTD and Mendelian randomization method, in which polygenic scores featured as genetic instruments. Bates et al. (2018) and Kong et al. (2018) proposed the use of polygenic scores based on transmitted and non-transmitted alleles specifically to detect genotype-environment covariance in the genetically informative trio design of parents and offspring. This method may obviously be applied in the extended twin design, including twins and their parents. To the best of our knowledge, the incorporation of polygenic scores in the twin design specifically to estimate A-E and A-C covariance has not been considered before.

The outline of this paper is as follows. First, we present the twin model, and the twin model extended with polygenic scores. Second, given the model for PGS in MZ and DZ twin pairs, we address the issues of parameter identification and power using exact data simulation. Third, we present the results of our power study. We conclude with a discussion.

## The twin model with polygenic scores

Let Y denote the phenotypic outcome variable of interest, and let GV_k_ denote the k-th genetic variant (GV) contributing to the variance of Y, where k = 1 … K, and K is the number of GVs contributing to individual differences in Y. We limit ourselves to an additive model, i.e., the GVs are additively coded (e.g., in the case of an di-allelic GV, 0, 1, or 2). The question how much of the variance in Y is explained by the GV_k_(k = 1,…,K) is addressed in the following regression model:

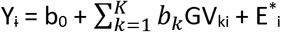

where b_0_ is the intercept, b_k_ is the k-th regression coefficient, subscript i denotes person, and E^*^ is the residual. Given a set of L of measured GVs, which are associated with the phenotypic Y, and the complementary set M unmeasured GVs (L + M = K), we have

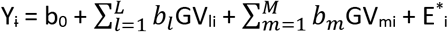

Given estimates of the regression coefficients (b_l_) obtained in an independent GWA study, the polygenic risk score equals 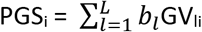 (e.g. Purcell et al. 2009; Dudbridge, 2013). The set L need not necessarily include GVs, which pass the genome-wide α-level. To ease presentation, we assume GVs are in linkage equilibrium (uncorrelated), so that the decomposition of phenotypic variance is

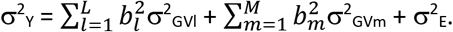

In the classical twin design, the ACE decomposition is:

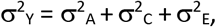

where we have partitioned the residual environmental variance σ^2^_E*_ into σ^2^_C_ and σ^2^_E_. Given the presence of the polygenic scores, we can decompose the additive genetic variance σ^2^_A_ into the observed component σ^2^_AL_ and the latent components σ^2^_AL_:

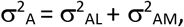

where 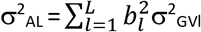 (the variance of the polygenic scores, i.e., σ^2^_AL_ = σ^2^_PGS_), and 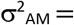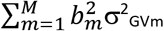. The information in the classical univariate twin model, i.e., the 2×2 MZ and DZ covariance matrices, is sufficient to obtain unique estimates of the variance components σ^2^_A_, σ^2^_C_, and σ^2^_E_. Figure 1 depicts the standard (ACE) twin model.

**Figure 1:**
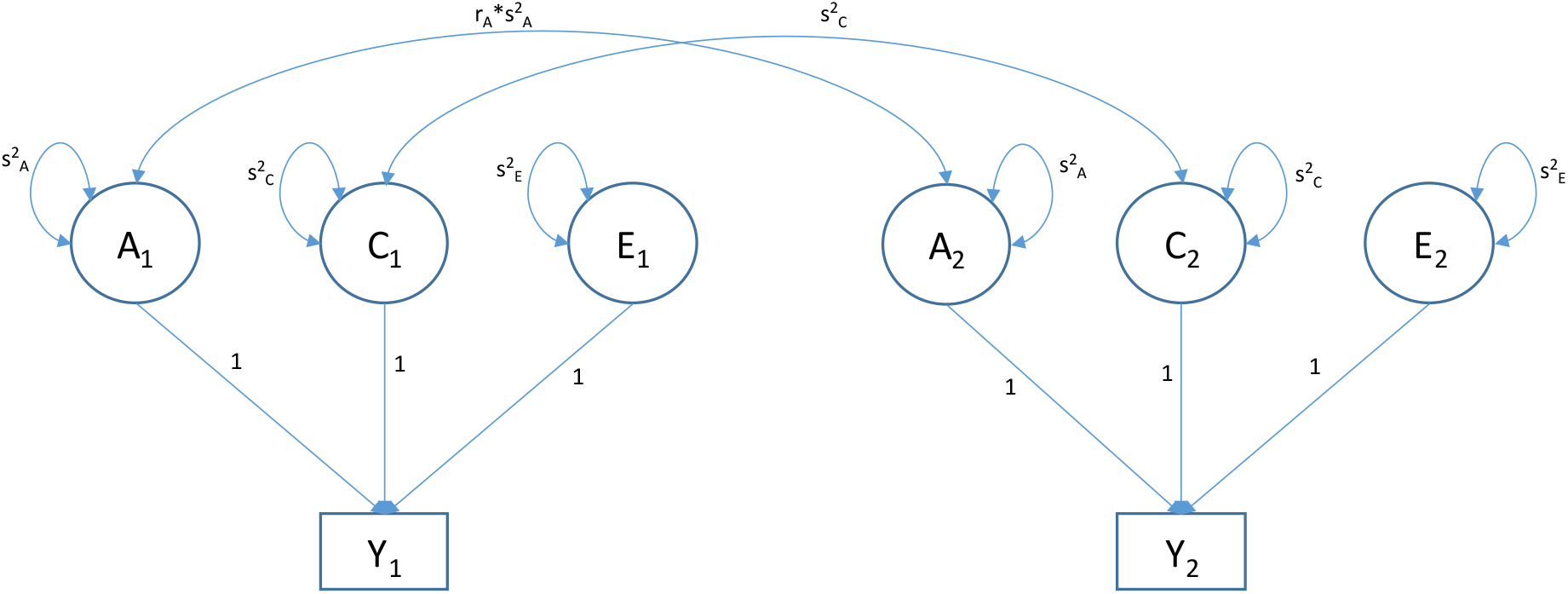
standard ACE twin model (r_A_ = 1 in MZs and r_A_= .5 in DZs)

The twin model including the polygenic scores is depicted in Figures 2 and 3 (to avoid clutter). Figure 2 depicts the model with A-C covariance parameter σ_AC_, and Figure 3 depicts the model with the A-E covariance parameter σ_AE_. The parameter R^2^ is the proportion of additive genetic variance attributable to 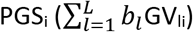, i.e., σ^2^_AL_/σ^2^_A_ or σ^2^_PGS_/σ^2^_A_. Limiting the model presentation to the covariance structure, we note that the standard ACE model (Figure 1) has three parameters (variance components σ^2^_A_, σ^2^, and σ^2^_E_), whereas the extended twin model has 6 parameters, i.e., the three variance components σ^2^_A_, σ^2^_C_, and σ^2^_E_, two covariance terms σ_AE_ and σ_AC_, and the parameter R^2^. The role of these parameters are shown in Table 1. Below we define effect sizes in terms of correlations, i.e., r_AC_ = σ_AC_ / (σ_A_*σ_C_) and r_AE_ = σ_AE_ / (σ_A_*σ_E_).

**Table 1.**
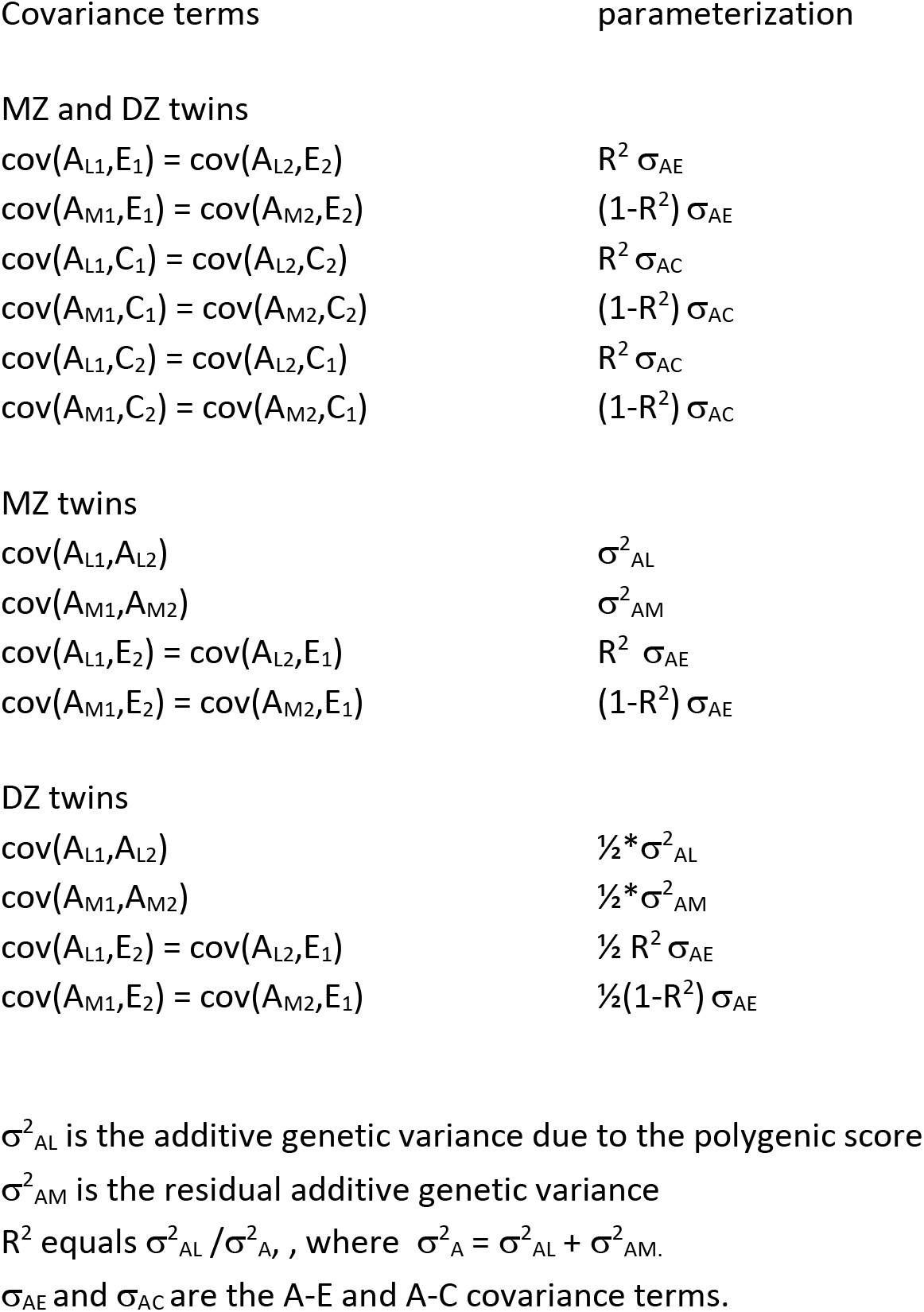
Covariance terms (see Figures 2 and 3).

| Covariance terms | parameterization |
| --- | --- |
| MZ and DZ twins |  |
| $\text{cov}(A_{L1}, E_1) = \text{cov}(A_{L2}, E_2)$ | $R^2 \sigma_{AE}$ |
| $\text{cov}(A_{M1}, E_1) = \text{cov}(A_{M2}, E_2)$ | $(1-R^2) \sigma_{AE}$ |
| $\text{cov}(A_{L1}, C_1) = \text{cov}(A_{L2}, C_2)$ | $R^2 \sigma_{AC}$ |
| $\text{cov}(A_{M1}, C_1) = \text{cov}(A_{M2}, C_2)$ | $(1-R^2) \sigma_{AC}$ |
| $\text{cov}(A_{L1}, C_2) = \text{cov}(A_{L2}, C_1)$ | $R^2 \sigma_{AC}$ |
| $\text{cov}(A_{M1}, C_2) = \text{cov}(A_{M2}, C_1)$ | $(1-R^2) \sigma_{AC}$ |
| MZ twins |  |
| $\text{cov}(A_{L1}, A_{L2})$ | $\sigma_{AL}^2$ |
| $\text{cov}(A_{M1}, A_{M2})$ | $\sigma_{AM}^2$ |
| $\text{cov}(A_{L1}, E_2) = \text{cov}(A_{L2}, E_1)$ | $R^2 \sigma_{AE}$ |
| $\text{cov}(A_{M1}, E_2) = \text{cov}(A_{M2}, E_1)$ | $(1-R^2) \sigma_{AE}$ |
| DZ twins |  |
| $\text{cov}(A_{L1}, A_{L2})$ | $\frac{1}{2} * \sigma_{AL}^2$ |
| $\text{cov}(A_{M1}, A_{M2})$ | $\frac{1}{2} * \sigma_{AM}^2$ |
| $\text{cov}(A_{L1}, E_2) = \text{cov}(A_{L2}, E_1)$ | $\frac{1}{2} R^2 \sigma_{AE}$ |
| $\text{cov}(A_{M1}, E_2) = \text{cov}(A_{M2}, E_1)$ | $\frac{1}{2} (1-R^2) \sigma_{AE}$ |

$\sigma_{AL}^2$ is the additive genetic variance due to the polygenic score
$\sigma_{AM}^2$ is the residual additive genetic variance
$R^2$ equals $\sigma_{AL}^2 / \sigma_A^2$ , , where $\sigma_A^2 = \sigma_{AL}^2 + \sigma_{AM}^2$ .
$\sigma_{AE}$ and $\sigma_{AC}$ are the A-E and A-C covariance terms.

**Figure 2:**
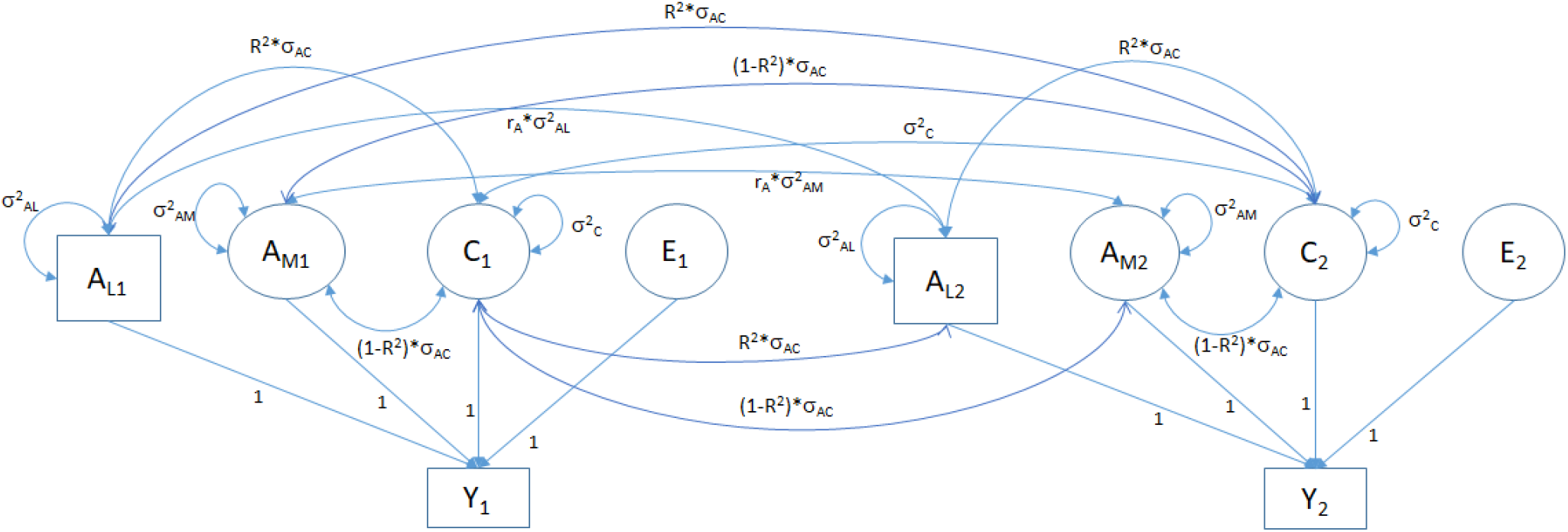
ACE twin model with polygenic scores (A_L1_, A_L2_), including A-E covariance (σ_AE_)

**Figure 3:**
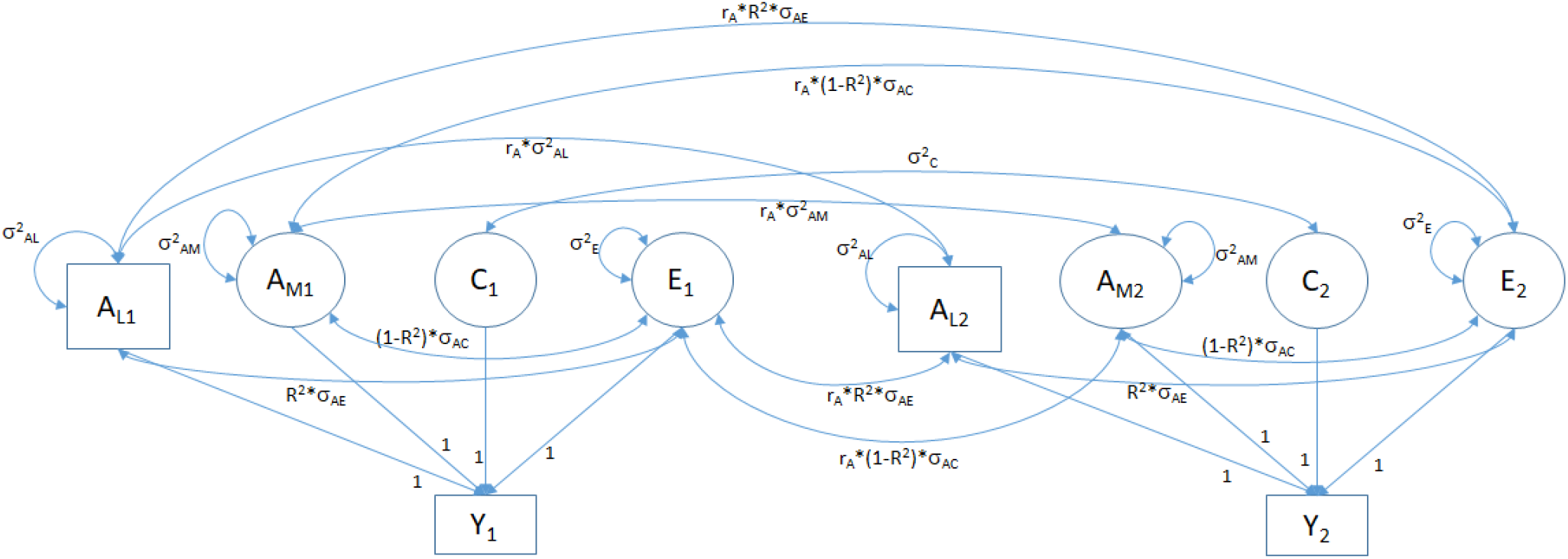
ACE twin model with polygenic scores (A_L1_, A_L2_), including A-C covariance (σ_AC_)

## Identification and resolution

The model is identified if the observed data provide sufficient information to obtain unique estimates of the unknown parameters when the likelihood of the data is maximized. In the present case, the observed information are the 3×3 MZ and the 4×4 DZ covariance matrices, containing the variances of, and the covariance among, the phenotype and the polygenic scores obtained in the twins. The MZ covariance matrix is 3×3, as MZ twins, being genetically identical, have identical polygenic scores.

As the parameterization of the phenotypic means is irrelevant to the issue of the identification of the covariance structure model, we simply estimate each mean with a separate free parameter, and do not discuss them further. Let the vector θ contain the parameters of the covariance part of the model:

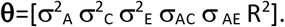

The vector θ does not include σ^2^_AL_ and σ^2^_AK_ explicitly, because these equal R^2^*σ^2^_A_ and (1-R^2^)*σ^2^_A_, respectively. Local identification implies that different points in the vicinity of a given point in the admissible 6 dimensional parameter space θ are associated with different expected covariance matrices Σ_MZ_(θ) and Σ_DZ_(θ) (Bollen and Bauldry, 2010; Bekker et al 1994). That is, we require that a change in the numerical value of one or more of the components in θ give rise to a change in the matrices matrices Σ_MZ_(θ) and Σ_DZ_(θ). We evaluated local identification analytically using symbolic algebra in Maple (Heck, 2003). The local identification check centers on the rank of the Jacobian matrix (Bekker et al 1994). The Jacobian matrix contains the first-order derivatives of the elements in the matrices Σ_MZ_(θ) and Σ_DZ_(θ) with respect to the parameters in θ. If the Jacobian is full column rank (as established using Maple), the model is locally identified. The advantage of this method is that it is necessary and sufficient; it does not depend on arbitrary numerical values of the parameters to establish local identification. We refer to Derks et al. (2006) and Minică et al. (2018) for other applications of this method in the context of twin modeling.

Having investigated local identification analytically, we proceeded to address the question of resolution by considering the statistical power to detect the parameters of interest. The issue of power is important, as formal local identification means that we can obtain unique parameter estimates, but tells us nothing about their precision. Specifically, the model may be formally identified, but empirically underidentified for certain sets of parameter values. We addressed the issue of power by conducting power analyses using exact data simulation (van de Sluis et al. 2008). We considered three parameter settings relating to the variance components, and combined them with various parameter R^2^ values and covariance parameters r_AC_ and r_AE_ (see Tables 2-4). We chose the values R^2^ = .15 and R^2^ =.05, and set the values of r_AC_ and r_AE_ to equal 0, .15, or .25. The percentage of phenotypic variance due to the polygenic score is included in the Tables (2–5). For instance, in a model with σ^2^_A_ =.333, σ^2^_C_ =.333, and σ^2^_E_ =.333 (see Table 2A,B), the polygenic score variance components equal .333/.15=.05 or .333/.05=0.0167. As the total phenotypic variance equals 1.333, the polygenic scores account for equals 100*(.05/1.333) = ~3.75% (R^2^=.15) or 100*(.0167/1.333) (R^2^=.05). The percentage of phenotypic variance explained polygenic score necessarily varies as a function of the parameter settings. That is, given a fixed value of R^2^, this percentage will vary as a function of the other parameter values. We test the A-C covariance and A-E covariance in isolation and simultaneously in 36 additional parameter settings (see Tables 2, 3, and 4). As the effects of C are generally absent in adults (e.g., in intelligence, personality, psychopathology), we also considered the detection of σ_AE_ ≠ 0 in an AE model (i.e., with σ^2^_C_=0 and r_AC_=0). The parameter settings are shown in Table 5.

**Table 2A.** σ^2^_A_ =.333, σ^2^_C_ =.333, σ^2^_E_ =.333

| set | R<sup>2</sup> | r<sub>AC</sub> | r<sub>AE</sub> | % | r<sub>AC</sub>=0<br>-<br>1 df<br>power | r<sub>EC</sub>=0<br>-<br>1df<br>power | r<sub>AC</sub>=0 & r<sub>AE</sub>=0<br>r<sub>AE</sub>=0<br>2 df<br>power | r<sub>AC</sub>=0 \| r<sub>AE</sub>=0<br>r<sub>AE</sub>=0<br>1 df<br>power | r<sub>AE</sub>=0 \| r<sub>AC</sub>=0<br>r<sub>AC</sub>=0<br>1df<br>power |
| --- | --- | --- | --- | --- | --- | --- | --- | --- | --- |
| 2.1 | .05 | .15 | 0 | 1.52 | .092 | - | .134 | .092 | - |
| 2.2 | .15 | .15 | 0 | 4.55 | .183 | - | .339 | .183 | - |
| 2.3 | .05 | .25 | 0 | 1.43 | .165 | - | .285 | .165 | - |
| 2.4 | .15 | .25 | 0 | 4.29 | .410 | - | **.733** | .410 | - |
| 2.5 | .05 | 0 | .15 | 1.52 | - | .093 | .142 | - | .183 |
| 2.6 | .15 | 0 | .15 | 4.55 | - | .187 | .366 | - | .461 |
| 2.7 | .05 | 0 | .25 | 1.43 | - | .164 | .311 | - | .398 |
| 2.8 | .15 | 0 | .25 | 4.29 | - | .409 | **.776** | - | **.855** |
| 2.9 | .05 | .15 | .15 | 1.39 | .088 | .092 | .372 | - | - |
| 2.10 | .15 | .15 | .15 | 4.17 | .173 | .186 | **.858** | - | - |
| 2.11 | .05 | .25 | .25 | 1.25 | .150 | .164 | **.761** | - | - |
| 2.12 | .15 | .25 | .25 | 3.75 | .369 | .408 | **.999** | - | - |

$\alpha = 0.05$ ; df = degrees of freedom; % = percentage of phenotypic variance due to the polygenic score; Nmz=Ndz=1000. $r_{AE}=0 \mid r_{AC}=0$ denotes the test of $r_{AE}=0$ , where $r_{AC}$ is fixed to zero (consistent with the model). ( $r_{AC}=0 \mid r_{AE}=0$ defined in the same way).

**Table 2B.** σ^2^_A_ =.333, σ ^2^_C_ =.333, σ^2^_E_ =.333.

| set | R<sup>2</sup> | r<sub>AC</sub> | r<sub>AE</sub> | % | r<sub>AC</sub>=0<br>-<br>1 df<br>power | r<sub>EC</sub>=0<br>-<br>1df<br>power | r<sub>AC</sub>=0 & r<sub>AE</sub>=0<br>r<sub>AE</sub>=0<br>2 df<br>power | r<sub>AC</sub>=0 \| r<sub>AE</sub>=0<br>r<sub>AE</sub>=0<br>1 df<br>power | r<sub>AE</sub>=0 \| r<sub>AC</sub>=0<br>r<sub>AC</sub>=0<br>1df<br>power |
| --- | --- | --- | --- | --- | --- | --- | --- | --- | --- |
| 2.1 | .05 | .15 | 0 | 1.52 | .151 | - | .134 | .151 | - |
| 2.2 | .15 | .15 | 0 | 4.55 | .293 | - | .339 | .293 | - |
| 2.3 | .05 | .25 | 0 | 1.43 | .317 | - | .285 | .632 | - |
| 2.4 | .15 | .25 | 0 | 4.29 | .632 | - | **.733** | .632 | - |
| 2.5 | .05 | 0 | .15 | 1.52 | - | .454 | .429 | - | .531 |
| 2.6 | .15 | 0 | .15 | 4.55 | - | .487 | .388 | - | .692 |
| 2.7 | .05 | 0 | .25 | 1.43 | - | **.848** | **.848** | - | **.909** |
| 2.8 | .15 | 0 | .25 | 4.29 | - | **.878** | **.956** | - | **.979** |
| 2.9 | .05 | .15 | .15 | 1.39 | .141 | .450 | .624 | - | - |
| 2.10 | .15 | .15 | .15 | 4.17 | .268 | .483 | **.923** | - | - |
| 2.11 | .05 | .25 | .25 | 1.25 | .280 | **.842** | **.963** | - | - |
| 2.12 | .15 | .25 | .25 | 3.75 | .561 | **.873** | **1.00** | - | - |

$\alpha = 0.05$ ; df = degrees of freedom; % = percentage of phenotypic variance due to the polygenic score; Nmz=Ndz=1000, R<sup>2</sup> fixed to true value. $r_{AE}=0 \mid r_{AC}=0$ denotes the test of $r_{AE}=0$ , where $r_{AC}$ is fixed to zero (consistent with the model). ( $r_{AC}=0 \mid r_{AE}=0$ defined in the same way).

**Table 3A.** σ^2^_A_ = .5, σ^2^_C_= .25, σ^2^_E_=.25.

| set | $R^2$ | $r_{AC}$ | $r_{AE}$ | % | $r_{AC}=0$<br>-<br>1 df<br>power | $r_{EC}=0$<br>-<br>1df<br>power | $r_{AC}=0$ & $r_{AE}=0$<br>$r_{AC}=0$ $r_{AE}=0$<br>2 df<br>power | $r_{AC}=0$ \| $r_{AE}=0$<br>$r_{AC}=0$ $r_{AE}=0$<br>1 df<br>power | $r_{AE}=0$ \| $r_{AC}=0$<br>$r_{AE}=0$ $r_{AC}=0$<br>1df<br>power |
| --- | --- | --- | --- | --- | --- | --- | --- | --- | --- |
| 3.1 | .05 | .15 | 0 | 2.26 | .081 | - | .110 | .081 | - |
| 3.2 | .15 | .15 | 0 | 6.78 | .151 | - | .262 | .151 | - |
| 3.3 | .05 | .25 | 0 | 2.12 | .136 | - | .219 | .136 | - |
| 3.4 | .15 | .25 | 0 | 6.37 | .328 | - | .597 | .328 | - |
| 3.5 | .05 | 0 | .15 | 2.26 | - | .082 | .117 | - | .148 |
| 3.6 | .15 | 0 | .15 | 6.78 | - | .154 | .284 | - | .366 |
| 3.7 | .05 | 0 | .25 | 2.12 | - | .135 | .238 | - | .309 |
| 3.8 | .15 | 0 | .25 | 6.37 | - | .326 | .644 | - | **.743** |
| 3.9 | .05 | .15 | .15 | 2.06 | .078 | .082 | .283 | - | - |
| 3.10 | .15 | .15 | .15 | 6.19 | .142 | .153 | **.736** | - | - |
| 3.11 | .05 | .25 | .25 | 1.85 | .124 | .134 | .617 | - | - |
| 3.12 | .15 | .25 | .25 | 5.54 | .292 | .325 | **.988** | - | - |

$\alpha = 0.05$ ; df = degrees of freedom; % = percentage of phenotypic variance due to the polygenic score; $Nmz=Ndz=1000$ . $r_{AE}=0$ | $r_{AC}=0$ denotes the test of $r_{AE}=0$ , where $r_{AC}$ is fixed to zero (consistent with the model). ( $r_{AC}=0$ | $r_{AE}=0$ defined in the same way).

**Table 3B.** σ ^2^_A_ = .5, σ^2^_C_= .25, σ^2^_E_=.25.

| set | $R^2$ | $r_{AC}$ | $r_{AE}$ | % | $r_{AC}=0$<br>-<br>1 df<br>power | $r_{EC}=0$<br>-<br>1df<br>power | $r_{AC}=0$ & $r_{AE}=0$<br>$r_{AC}=0$ $r_{AE}=0$<br>2 df<br>power | $r_{AC}=0$ \| $r_{AE}=0$<br>$r_{AC}=0$ $r_{AE}=0$<br>1 df<br>power | $r_{AE}=0$ \| $r_{AC}=0$<br>$r_{AE}=0$ $r_{AC}=0$<br>1df<br>power |
| --- | --- | --- | --- | --- | --- | --- | --- | --- | --- |
| 3.1 | .05 | .15 | 0 | 2.26 | .129 | - | .110 | .129 | - |
| 3.2 | .15 | .15 | 0 | 6.78 | .259 | - | .262 | .259 | - |
| 3.3 | .05 | .25 | 0 | 2.12 | .260 | - | .219 | .260 | - |
| 3.4 | .15 | .25 | 0 | 6.37 | .566 | - | .597 | .566 | - |
| 3.5 | .05 | 0 | .15 | 2.26 | - | .526 | .474 | - | .578 |
| 3.6 | .15 | 0 | .15 | 6.78 | - | .539 | .583 | - | .687 |
| 3.7 | .05 | 0 | .25 | 2.12 | - | **.907** | **.888** | - | **.937** |
| 3.8 | .15 | 0 | .25 | 6.37 | - | **.916** | **.953** | - | **.977** |
| 3.9 | .05 | .15 | .15 | 2.06 | .121 | .523 | .613 | - | - |
| 3.10 | .15 | .15 | .15 | 6.19 | .237 | .536 | **.874** | - | - |
| 3.11 | .05 | .25 | .25 | 1.85 | .229 | **.903** | **.961** | - | - |
| 3.12 | .15 | .25 | .25 | 5.54 | .496 | **.912** | **.999** | - | - |

$\alpha = 0.05$ ; df = degrees of freedom; % = percentage of phenotypic variance due to the polygenic score; $Nmz=Ndz=1000$ , $R^2$ fixed to true value. $r_{AE}=0$ | $r_{AC}=0$ denotes the test of $r_{AE}=0$ , where $r_{AC}$ is fixed to zero (consistent with the model). ( $r_{AC}=0$ | $r_{AE}=0$ defined in the same way).

**Table 4A.** σ^2^_A_ = .25, σ^2^_C_ = .5, σ^2^_E_=. 25.

| set | $R^2$ | $r_{AC}$ | $r_{AE}$ | % | $r_{AC}=0$<br>-<br>1 df<br>power | $r_{EC}=0$<br>-<br>1df<br>power | $r_{AC}=0$ & $r_{AE}=0$<br>$r_{AC}=0$ \| $r_{AE}=0$<br>2 df<br>power | $r_{AC}=0$ \| $r_{AE}=0$<br>1 df<br>power | $r_{AE}=0$ \| $r_{AC}=0$<br>1df<br>power |
| --- | --- | --- | --- | --- | --- | --- | --- | --- | --- |
| 4.1 | .05 | .15 | 0 | 1.13 | .122 | - | .170 | .122 | - |
| 4.2 | .15 | .15 | 0 | 3.39 | .281 | - | .451 | .281 | - |
| 4.3 | .05 | .25 | 0 | 1.06 | .250 | - | .384 | .250 | - |
| 4.4 | .15 | .25 | 0 | 3.19 | .620 | - | **.868** | .620 | - |
| 4.5 | .05 | 0 | .15 | 1.16 | - | .092 | .123 | - | .157 |
| 4.6 | .15 | 0 | .15 | 3.49 | - | .185 | .301 | - | .386 |
| 4.7 | .05 | 0 | .25 | 1.11 | - | .163 | .261 | - | .337 |
| 4.8 | .15 | 0 | .25 | 3.33 | - | .405 | .681 | - | **.776** |
| 4.9 | .05 | .15 | .15 | 1.06 | .117 | .092 | .394 | - | - |
| 4.10 | .15 | .15 | .15 | 3.18 | .265 | .185 | **.877** | - | - |
| 4.11 | .05 | .25 | .25 | 0.96 | .226 | .162 | **.794** | - | - |
| 4.12 | .15 | .25 | .25 | 2.88 | .573 | .404 | **.999** | - | - |

$\alpha = 0.05$ ; df = degrees of freedom; % = percentage of phenotypic variance due to the polygenic score; $Nmz=Ndz=1000$ . $r_{AE}=0$ | $r_{AC}=0$ denotes the test of $r_{AE}=0$ , where $r_{AC}$ is fixed to zero (consistent with the model). ( $r_{AC}=0$ | $r_{AE}=0$ defined in the same way).

**Table 4B.** σ^2^_A_ = .25, σ^2^_C_ = .5, σ^2^_E_=. 25.

| set | $R^2$ | $r_{AC}$ | $r_{AE}$ | % | $r_{AC}=0$<br>-<br>1 df<br>power | $r_{EC}=0$<br>-<br>1df<br>power | $r_{AC}=0$ & $r_{AE}=0$<br>$r_{AC}=0$ \| $r_{AE}=0$<br>2 df<br>power | $r_{AC}=0$ \| $r_{AE}=0$<br>1 df<br>power | $r_{AE}=0$ \| $r_{AC}=0$<br>1df<br>power |
| --- | --- | --- | --- | --- | --- | --- | --- | --- | --- |
| 4.1 | .05 | .15 | 0 | 1.13 | .199 | - | .170 | .199 | - |
| 4.2 | .15 | .15 | 0 | 3.39 | .420 | - | .451 | .420 | - |
| 4.3 | .05 | .25 | 0 | 1.06 | .435 | - | .384 | .435 | - |
| 4.4 | .15 | .25 | 0 | 3.19 | **.815** | - | **.868** | **.815** | - |
| 4.5 | .05 | 0 | .15 | 1.16 | - | .445 | .402 | - | .501 |
| 4.6 | .15 | 0 | .15 | 3.49 | - | .478 | .528 | - | .633 |
| 4.7 | .05 | 0 | .25 | 1.11 | - | **.839** | **.821** | - | **.889** |
| 4.8 | .15 | 0 | .25 | 3.33 | - | **.871** | **.928** | - | **.962** |
| 4.9 | .05 | .15 | .15 | 1.06 | .188 | .442 | .636 | - | - |
| 4.10 | .15 | .15 | .15 | 3.18 | .392 | .475 | **.933** | - | - |
| 4.11 | .05 | .25 | .25 | 0.96 | **.396** | **.836** | **.968** | - | - |
| 4.12 | .15 | .25 | .25 | 2.88 | **.762** | **.868** | **1.00** | - | - |

$\alpha = 0.05$ ; df = degrees of freedom; % = percentage of phenotypic variance due to the polygenic score; $Nmz=Ndz=1000$ , $R^2$ fixed to true value. $r_{AE}=0$ | $r_{AC}=0$ denotes the test of $r_{AE}=0$ , where $r_{AC}$ is fixed to zero (consistent with the model). ( $r_{AC}=0$ | $r_{AE}=0$ defined in the same way).

**Table 5.**
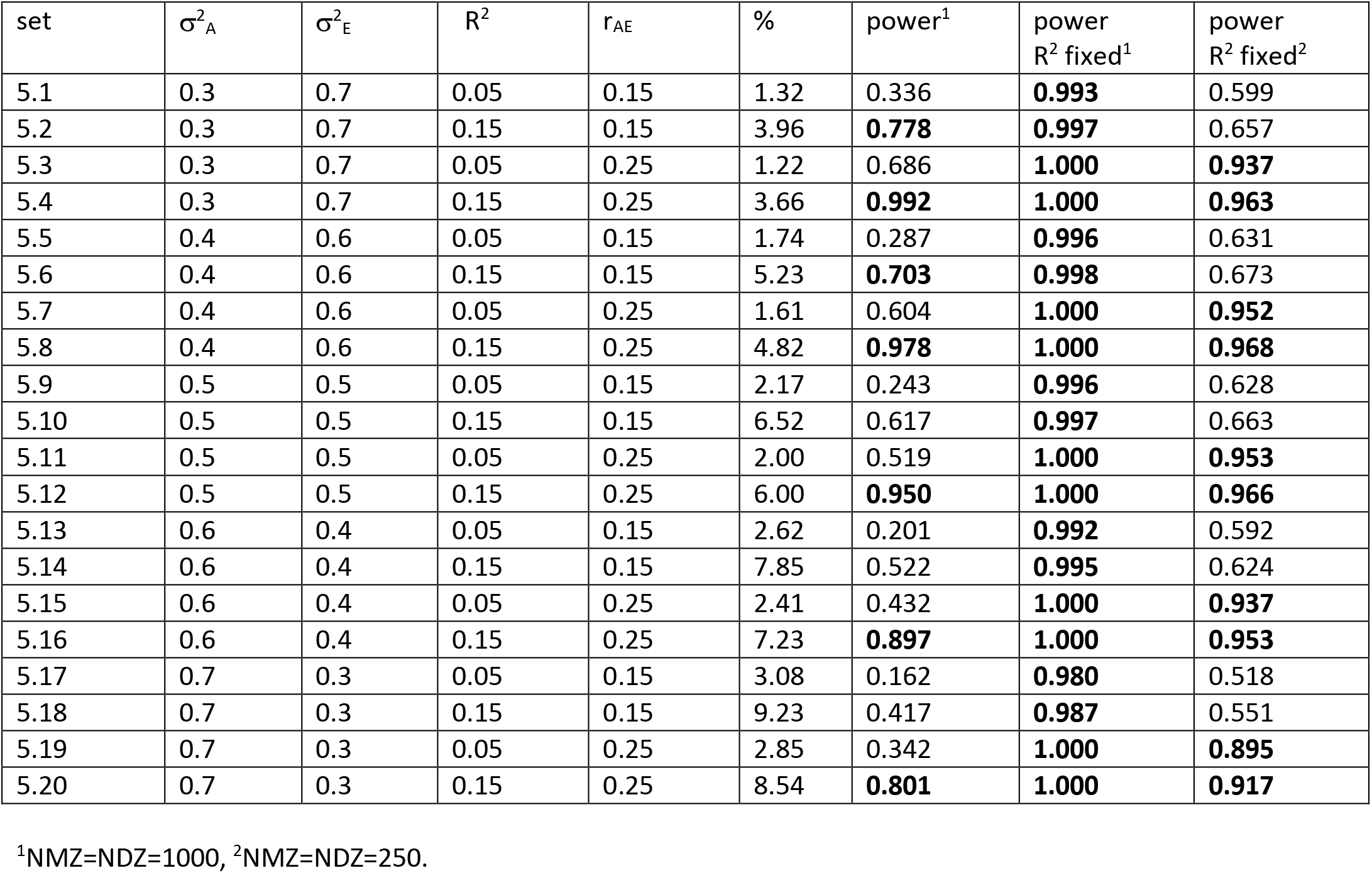
Parameter settings, power to reject r_AE_=0 (1 df likelihood ratio test), and the non-centrality parameter values. (α = 0.05). Column 7 and 8 “power R^2^ fixed” is the power to reject r_AE_=0, given R^2^ fixed to the true value, shown in column 4. “%” is the percentage of phenotypic variance due to the polygnic score.

In all cases, we set the sample sizes equal to Nmz=1000 and Ndz=1000 (in total 2000 pairs), and we adopted an α of 0.05. We used normal theory maximum likelihood estimation throughout, and based our power calculations on the non-centrality parameter associated with the chi-square distribution (Martin, et al. 1978). The power analyses were done using OpenMx (Boker et al 2011; Neale et al 2016) in the R program (R core team, 2018). The OpenMx script is available (supplemental material), so that readers may calculate the power in other settings.

## Results

The analytical check using Maple demonstrated that the model is identified, on the basis of the 3×3 MZ and the 4×4 DZ phenotypic covariance matrices, Σ_MZ_(**θ**) and Σ_DZ_(**θ**). That is, we can, at least in principle, obtain unique estimates of **θ**=[σ^2^_A_ σ^2^_C_ σ^2^_E_ σ_AC_ σ _AE_ R^2^]. Tables 2A, 3A, and 4A shows the power given our parameter settings (associated non-centrality parameters are given in the supplemental material). The power varies considerably in Table 2A (σ^2^_A_ =.333, σ^2^_C_ =.333, σ^2^_E_ =.333), from .092 to .999. We observe power greater than the (arbitrary) value of .70 is observed in 6 cases. Of these 6, 5 are associated with the omnibus (2 df) test r_AC_=0 <u>and</u> r_AE_ = 0. The 6th case involves the (1 df) test of r_AE_ = 0, given r_AC_ = 0 (consistent with the model in which r_AC_ = 0). Unsurprisingly, in all 6 cases, the R^2^ (i.e., σ^2^_PGS_/σ^2^_A_) equals .15, rather than .05. In these cases the polygenic scores explain between 3.7 and 4.29% of phenotypic variance. In Table 3A (σ^2^_A_ = .5, σ^2^_C_= .25, σ^2^_E_=.25), we see only three scenarios with power greater than .7. These again concern the 2df test (r_AE_=0 & r_AC_=0) and a test of r_AE_ = 0, given r_AC_ = 0 (fixed to zero, consistent with the model) in the R^2^= .15 conditions. The results in Table 4A (σ^2^_A_ = .25, σ^2^_C_ = .5, σ^2^_E_=. 25) largely resemble those of Table 2A. As the number of cases with power >.70 are limited, we conclude that the method may be underpowered given Nmz = Ndz = 1000 and α=.05.

By way of exploration, we considered the power given a known value of the parameter R^2^, that is, we assume that we know how much the polygenic scores explain of the total additive genetic variance, for example from a large study in a comparable population. Tables 2B to 4A shows the power in the same parameter settings as given in Tables 2A to 4A, but with the parameter R^2^ fixed to its true value. The power is greatly improved by this measure. In Tables 2B, 3B, and 4B, we see 12, 11, and 16 cases, respectively, with power > .70. To obtain some insight into this finding, we calculated the correlations among the parameter estimates in case 2.12 (see Tables 2A and 2B), i.e., given σ^2^_A_ =.333, σ^2^_C_ =.333, σ^2^_E_ =.333, R^2^=.15, and r_AC_ = r_AE_ = .25. This correlation is available in the OpenMx output (i.e., the OpenMx vcov() function). We know from Table 2A and 2B that the power to reject r_AC_=0 is low at .369 (Table 2A, R^2^ estimated) or .561 (Table 2A, R^2^ fixed), and the power to reject r_AE_=0 is .408 (Table 2A, R^2^ estimated) or .873 (Table 2B, R^2^ fixed). The low power to reject r_AC_=0 (.369) is due in part to the relatively large correlation between the parameter estimates r_AC_ and r_AE_ (-.82), implying that the effect of fixing one parameter (to zero) will be compensated by the other. We note that the correlation between the estimates of the A and C variance terms (σ^2^_A_ and σ^2^_C_) is also large (-.90), as is well-known. Having fixed the R^2^ parameter, we find that the correlations among the parameters rAC and rAE are appreciably lower −0.61. Interestingly, the correlation between the estimates of σ^2^_A_ and σ^2^_C_ is only −.21, which implies that the inclusion of polygenic scores in the twin model will greatly increase the power to detect C variance. The power to detect C in the standard ACE model is known to be low in general (Visscher, Gordon, and Neale, 2008).

Table 5 shows the power to detect r_AE_ (in the absence of r_AC_) in the AE model. With R^2^ estimated, we see 7 cases (of the 20 considered) with power greater than .7. The largest power (.978) is associated with σ_A_^2^ = .4, σ_E_^2^ = .6. R^2^=.15 and r_AE_ = .25. We repeated the analyses with the R^2^ parameter fixed to its true value. The power in all cases exceeds .97 (see Table 5 column 7). To provide more insight into the variation in the power over the parameter settings considered, we repeated the analyses with Nmz=Ndz=250 (retaining α=0.05). Given Nmz=Ndz=250, we observe 10 cases with power greater than .7.

## Discussion

We addressed the question of whether incorporating polygenic scores into the classical twin model rendered the A-C and A-E correlations (or covariances) identified. Based on the analytical results relating to the column rank of the model's matrix of partial derivatives (Jacobian) matrix and given the estimation of correct numerical values in exact data simulation (van der Sluis et al, 2008), we established that both parameters, simultaneously estimated, are identified. The power to demonstrate that these correlations are greater than zero, in isolation (1df test) or together (2df test), given NMZ=NDZ=1000 and α=0.05, varied greatly as a function of the parameter settings. Overall, in view of the results in Tables 2A to 4A, we found the likelihood ratio tests to be underpowered in many cases. Fixing the R^2^ parameter to its true value helped greatly to remedy this lack of power (see Tables 2B to 4B). This solution is not too outlandish, as in practice, we often know the heritability of the phenotype of interest quite accurately (see Polderman et al. 2015), and the percentage of phenotypic variance explained by the polygenic scores. Based on that information, one can arrive at an approximate value of R^2^. Also, in practice, one may vary the fixed R^2^ values to gauge the sensitivity of the estimates of r_AC_ and r_AE_ to the value of R^2^.

We considered a polygenic score effect size that was expressed in terms of R^2^, the proportion of total additive genetic variance attributable to the polygenic score. The values .05 and .15 are arbitrary, but not overly large. As shown in the Tables (2-4), these values imply that the polygenic scores account for between 1.25 (R^2^=.05) and 4.55 (Table 2A), between 1.85 (R^2^=.05) and 6.78 (Table 3A), and between 1.06 (R^2^=.05) and 3.49 % (Table 4A) of the phenotypic variance. If polygenic scores of the phenotype of interest at present do not reach these levels, they will likely do so in the future, given the results of ever larger genome wide association meta-analyses.

We have focused on the test of r_AC_ and r_AC_ in the basic univariate twin model. Clearly the incorporation of polygenic scores may facilitate detection and modeling of r_AC_ and r_AE_ in other models. From a developmental point of view, a longitudinal study of cognitive ability would benefit greatly by the incorporation of polygenic scores to determine whether age-related changes in the A and C variance components of intelligence (e.g., the decrease in C variance) are dependent on age-related changes in r_AC_ (Haworth et al. 2010, Tucker-Drob & Bates, 2016). While we have focused on covariance, we note that polygenic scores have wider applications in the study of genotype-environment interplay. Polygenic scores are already being used to study interaction (e.g., Colodro-Conde et al 2018; Peyrot, et al. 2014). Assigning polygenic scores the role of moderator in the twin moderation model (Purcell, 2002) will facilitate the study of AxE and AxC interaction, allowing for the developmentally important distinction of AxC and AxE. By comparison, the method proposed by Molenaar et al (2012) is computationally more complex and likely to be less powerful, as it is based on the phenotypic twin data without polygenic scores.

In conclusion, the incorporation of polygenic scores in the twin model provides new possibilities to study interplay, as manifested in A-C and A-E covariance. We found that the present approach was often underpowered, given Nmz = Ndz =1000 and α=0.05, if the parameter R^2^ was estimated along with the other parameters in the model. To improve power, we proposed to fix R^2^ to a well-informed value (based on knowledge of the variance explained by the polygenic score). We suspect that in the coming years polygenic scores will gain statistical power, as their effect sizes (in terms of the amount of additive genetic variance explained) will increase, and their standard errors will decrease (due to larger sample sizes), so that ultimately this "fix" will not be necessary.

## Acknowledgements

CVD acknowledges NWO top-talent grant 406-12-124 (awarded to Janneke de Kort); DIB acknowledges KNAW Academy Professor Award (PAH/6635); MCN and CCM were supported by the National Institute on Drug Abuse (DA-018673; PI: MCN).

